# Mosaic trisomy of chromosome 1q in human brain tissue associates with unilateral polymicrogyria, very early-onset focal epilepsy, and severe developmental delay

**DOI:** 10.1101/2020.07.16.206490

**Authors:** Katja Kobow, Samir Jabari, Tom Pieper, Manfred Kudernatsch, Tilman Polster, Friedrich G Woermann, Thilo Kalbhenn, Hajo Hamer, Karl Rössler, Angelika Mühlebner, Wim GM Spliet, Martha Feucht, Yanghao Hou, Damian Stichel, Andrey Korshunov, Felix Sahm, Roland Coras, Ingmar Blümcke, Andreas von Deimling

**Author notes:** these authors contributed equally (Katja Kobow, Samir Jabari, Ingmar Blümcke, Andreas von Deimling). **Author for correspondence:** Katja Kobow, Universitätsklinikum Erlangen, Institute of Neuropathology, Schwabachanlage 6, 91054 Erlangen, Germany.

## Abstract

Polymicrogyria (PMG) is a developmental cortical malformation characterized by an excess of small and frustrane gyration and abnormal cortical lamination. PMG frequently associates with seizures. The molecular pathomechanisms underlying PMG development are not yet understood. About 40 genes have been associated with PMG, and small copy number variations have also been described in selected patients. We recently provided evidence that epilepsy-associated structural brain lesions can be classified based on genomic DNA methylation patterns. Here we analyzed 26 PMG patients employing array-based DNA-methylation profiling on formalin-fixed paraffin-embedded material. A series of 62 well-characterized non-PMG cortical malformations (focal cortical dysplasia type 2a/b and hemimegalencephaly), temporal lobe epilepsy, and non-epilepsy autopsy controls was used as reference cohort. Unsupervised dimensionality reduction and hierarchical cluster analysis of DNA methylation profiles showed that PMG formed a distinct DNA methylation class. Copy number profiling from DNA methylation data identified a uniform duplication spanning the entire long arm of chromosome 1 in 7 out of 26 PMG patients, which was verified by additional fluorescence in situ hybridization analysis. In respective cases about 50% of nuclei in the center of the PMG lesion were 1q triploid. No chromosomal imbalance was seen in adjacent, architecturally normal-appearing tissue indicating mosaicism. Clinically, PMG 1q patients presented with a unilateral frontal or hemispheric PMG without hemimegalencephaly, a severe form of intractable epilepsy with seizure onset in the first months of life, and severe developmental delay. Our results show that PMG can be classified among other structural brain lesions according to their DNA methylation profile. One subset of PMG with distinct clinical features exhibits a duplication of chromosomal arm 1q.

## Introduction

PMG is a malformation of cortical development (MCD) characterized by an excessive number of abnormally small and partly fused, and so called frustrane gyration together with abnormal cortical lamination [7]. PMG can be limited to a single gyrus, involving only part of one hemisphere, be bilateral and asymmetrical, bilateral and symmetrical, or diffuse [6, 54]. Clinical signs and symptoms are heterogeneous depending on how many and which brain regions are affected. They may range from mild intellectual disability, mobility and language problems to severe encephalopathy with intractable epilepsy. While PMG most often occurs as an isolated malformation, it can accompany several other brain malformations, e.g., microcephaly, megalencephaly, periventricular nodular heterotopia, FCD, agenesis of the corpus callosum and brainstem, or cerebellar abnormalities [60]. PMG is genetically heterogeneous with about 40 associated genes [31, 54, 62]. Small copy-number variants (CNVs) have also been associated with PMG, but only deletions in 1p36.3 and 22q11.2 are common, otherwise CNVs are rare [23, 56]. A causal gene has not been identified for any of the CNV loci. Among non-genetic causes of PMG hypoxia-ischemia, trauma or congenital infections mainly from cytomegalovirus have been reported [8, 37].

Microscopically, in PMG, the gyri are atypically organized with abnormalities of the physiological laminar cytoarchitectural structure, which may be unlayered or four-layered. In unlayered PMG, the molecular layer is continuous and does not follow convolutional profiles. Neurons have a radial distribution without any laminar organization. By contrast, the four-layered PMG shows a laminar structure composed of a molecular layer, an outer neuronal layer, a nerve fiber layer, and an inner neuronal layer. Occasionally, in the neuronal cell layers, granular as well as pyramidal neurons can be observed, resembling residuals of the normal six-layered cortical architecture [25]. The two histopathological PMG-subtypes do not necessarily have a distinct origin, as both may coexist in contiguous cortical regions [10].

Though PMG is frequently associated with epilepsy, mechanisms of epileptogenesis related to PMG are not fully understood. Experimental data obtained in the rat freeze lesion model indicate that functional abnormalities extend beyond the anatomical malformation [30, 39]. This corroborates observations in humans, revealing that the cortex surrounding the PMG is also involved in epileptogenesis.

Classification of MCD including PMG remains challenging in everyday clinical practice [40]. DNA methylation profiling can be used for molecular classification of seemingly morphological homogenous entities. It can further provide certain information on chromosomal imbalances, and inform about molecular pathways underlying disease development. This has been proven to variable extent in brain tumors and major subtypes of focal cortical dysplasia (FCD) [32, 36, 53, 57]. We hypothesized that the determination of DNA methylation signatures might help distinguish PMG from other related hemispheric and focal MCD and identify PMG subtypes with specific molecular and clinical features. In the present study, we, therefore, used llumina DNA Methylation 850K BeadChip Arrays to molecularly characterize 26 PMG and correlate molecular-genetic data with histomorphological and clinical phenotype.

## Material and Methods

### Study Subjects

We obtained written informed consent for molecular-genetic investigations and publication of the results for all participating patients. The Ethics Committee of the Medical Faculty of the Friedrich-Alexander-University (FAU) Erlangen-Nürnberg, Germany, approved this study within the framework of the EU project “DESIRE” (FP7, grant agreement 602531; AZ 92_14B). We reviewed clinical, imaging and histological data of individuals who underwent surgery for the treatment of their focal pharmacoresistent epilepsy and were diagnosed with unilateral PMG (n=26; mean age at surgery ± SEM = 12.5 ± 3.5 years; **Table 1**), hemimegalencephaly (HME, n=6; mean age ± SEM = 1.3 ± 0.2 years), FCD type 2 (n=36; mean age ± SEM = 15.8 ± 2.2 years), or temporal lobe epilepsy (TLE, n=15; mean age ± SEM = 37.0 ± 4.0 years; All TLE patients had a histopathological diagnosis of hippocampal sclerosis, but only apparently normal temporal neocortex was used in the present analysis.). All disease diagnosis was based on MRI and histology. Five non-epilepsy autopsy control cases with no known neurological history were also included in the study (mean age ± SEM = 30.4 ± 8.7 years; **Supplement Table 1**).

**Table 1:**
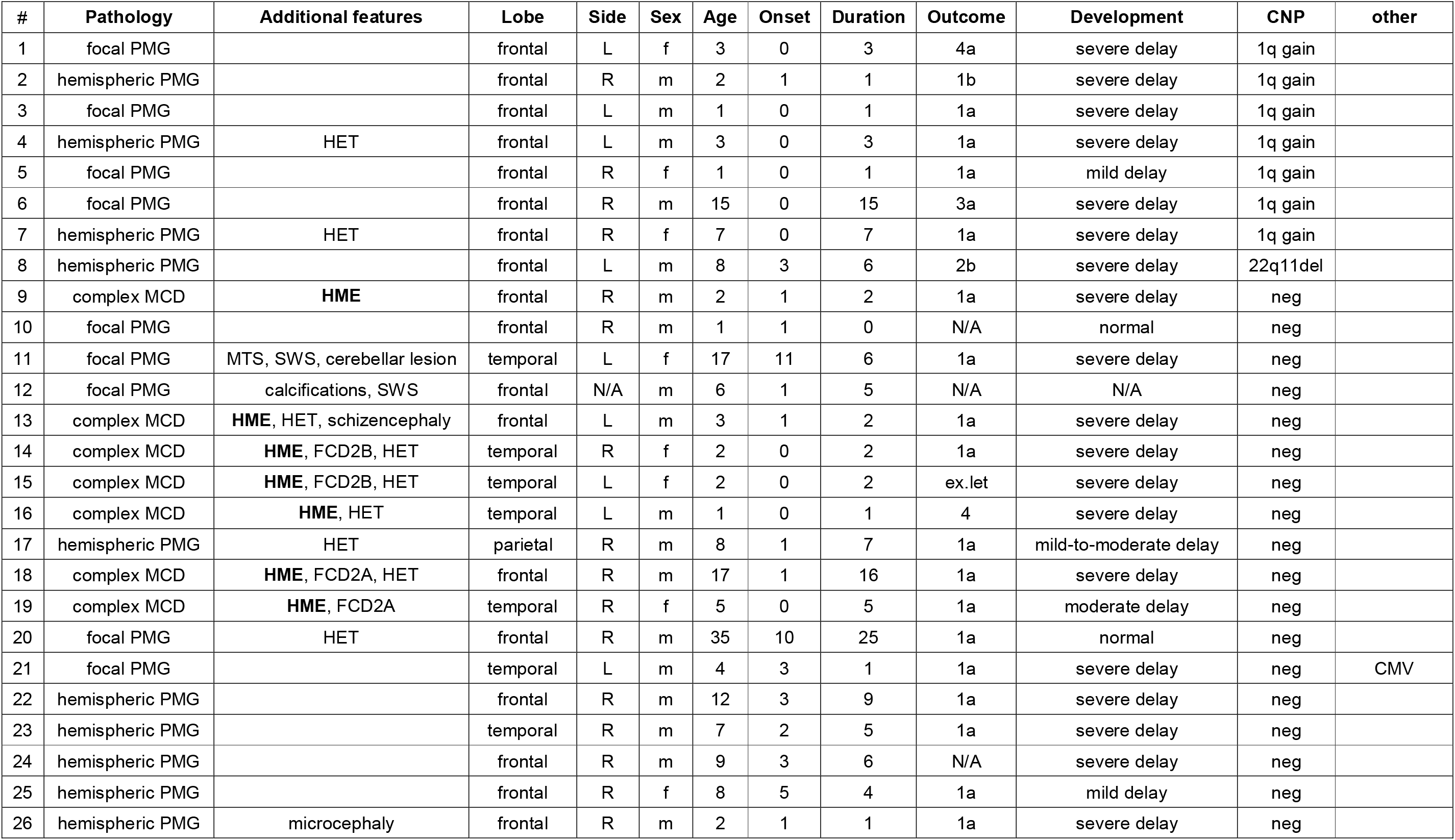
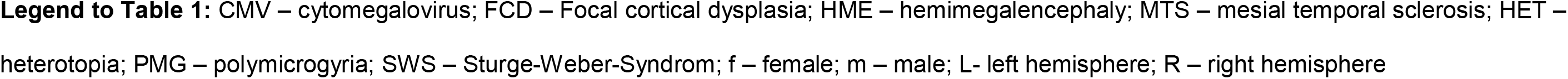
Clinical summary of PMG cases.

| # | Pathology | Additional features | Lobe | Side | Sex | Age | Onset | Duration | Outcome | Development | CNP | other |
| --- | --- | --- | --- | --- | --- | --- | --- | --- | --- | --- | --- | --- |
| 1 | focal PMG |  | frontal | L | f | 3 | 0 | 3 | 4a | severe delay | 1q gain |  |
| 2 | hemispheric PMG |  | frontal | R | m | 2 | 1 | 1 | 1b | severe delay | 1q gain |  |
| 3 | focal PMG |  | frontal | L | m | 1 | 0 | 1 | 1a | severe delay | 1q gain |  |
| 4 | hemispheric PMG | HET | frontal | L | m | 3 | 0 | 3 | 1a | severe delay | 1q gain |  |
| 5 | focal PMG |  | frontal | R | f | 1 | 0 | 1 | 1a | mild delay | 1q gain |  |
| 6 | focal PMG |  | frontal | R | m | 15 | 0 | 15 | 3a | severe delay | 1q gain |  |
| 7 | hemispheric PMG | HET | frontal | R | f | 7 | 0 | 7 | 1a | severe delay | 1q gain |  |
| 8 | hemispheric PMG |  | frontal | L | m | 8 | 3 | 6 | 2b | severe delay | 22q11del |  |
| 9 | complex MCD | HME | frontal | R | m | 2 | 1 | 2 | 1a | severe delay | neg |  |
| 10 | focal PMG |  | frontal | R | m | 1 | 1 | 0 | N/A | normal | neg |  |
| 11 | focal PMG | MTS, SWS, cerebellar lesion | temporal | L | f | 17 | 11 | 6 | 1a | severe delay | neg |  |
| 12 | focal PMG | calcifications, SWS | frontal | N/A | m | 6 | 1 | 5 | N/A | N/A | neg |  |
| 13 | complex MCD | HME, HET, schizencephaly | frontal | L | m | 3 | 1 | 2 | 1a | severe delay | neg |  |
| 14 | complex MCD | HME, FCD2B, HET | temporal | R | f | 2 | 0 | 2 | 1a | severe delay | neg |  |
| 15 | complex MCD | HME, FCD2B, HET | temporal | L | f | 2 | 0 | 2 | ex.let | severe delay | neg |  |
| 16 | complex MCD | HME, HET | temporal | L | m | 1 | 0 | 1 | 4 | severe delay | neg |  |
| 17 | hemispheric PMG | HET | parietal | R | m | 8 | 1 | 7 | 1a | mild-to-moderate delay | neg |  |
| 18 | complex MCD | HME, FCD2A, HET | frontal | R | m | 17 | 1 | 16 | 1a | severe delay | neg |  |
| 19 | complex MCD | HME, FCD2A | temporal | R | f | 5 | 0 | 5 | 1a | moderate delay | neg |  |
| 20 | focal PMG | HET | frontal | R | m | 35 | 10 | 25 | 1a | normal | neg |  |
| 21 | focal PMG |  | temporal | L | m | 4 | 3 | 1 | 1a | severe delay | neg | CMV |
| 22 | hemispheric PMG |  | frontal | R | m | 12 | 3 | 9 | 1a | severe delay | neg |  |
| 23 | hemispheric PMG |  | temporal | R | m | 7 | 2 | 5 | 1a | severe delay | neg |  |
| 24 | hemispheric PMG |  | frontal | R | m | 9 | 3 | 6 | N/A | severe delay | neg |  |
| 25 | hemispheric PMG |  | frontal | R | f | 8 | 5 | 4 | 1a | mild delay | neg |  |
| 26 | hemispheric PMG | microcephaly | frontal | R | m | 2 | 1 | 1 | 1a | severe delay | neg |  |

**Legend to Table 1:** CMV – cytomegalovirus; FCD – Focal cortical dysplasia; HME – hemimegalencephaly; MTS – mesial temporal sclerosis; HET – heterotopia; PMG – polymicrogyria; SWS – Sturge-Weber-Syndrom; f – female; m – male; L- left hemisphere; R – right hemisphere

### DNA and RNA extraction

A prototypical area within the center of the MCD lesion (neocortex) was identified on H&E slides and macrodissection performed by punch biopsy (pfm medical, Köln, Germany) or by hand. DNA and RNA were extracted from formalin-fixed paraffin-embedded (FFPE) tissue using the Maxwell 16 FFPE Plus LEV DNA Kit and Maxwell 16 LEV RNA FFPE Purification Kit (Promega, Madison, WI, USA), according to manufacturer’s instructions. DNA concentration was quantified using the Qubit dsDNA BR Assay kit (Invitrogen, Carlsbad, CA, USA).

### Genome-wide DNA methylation profiling and data pre-processing

Samples were analyzed using Illumina Infinium MethylationEPIC 850K BeadChip arrays, as described previously [16]. Briefly, DNA methylation data were generated at the Department of Neuropathology, Universitätsklinikum Heidelberg, Germany. Copy-number profile (CNP) analysis was assessed using R package “conumee” after an additional baseline correction (https://github.com/dstichel/conumee).

Differential DNA methylation analysis was performed with a self-customized Python wrapped cross R package pipeline, utilizing ‘Champ,’ ‘watermelon,’ ‘DNAMarray,’ and ‘minfi’ as R packages. ‘MethQC’ and ‘Pymethylarray’ were used as Python-based array analysis packages. Additional own implementations of lacking functionality were used where needed. All these packages were wrapped by using ‘rpy2’, an interface to R running embedded in a Python process. Methylation array data was read utilizing minfi‘s ‘read_metharray_sheet’ and ‘read_metharray_exp’ function as well as a self-modified parallelized version available from the ‘DNAMarray’ R-package. Data was stratified quantile normalized using the ‘minfi’ ‘preprocessQuantile’ function. Probes targeting sex chromosomes, probes containing single nucleotide polymorphisms (SNPs) not uniquely matching, as well as known cross-reactive probes (see [18]) were removed. Additionally, a rpy2 wrapped and modified version of the ‘reduce’ and ‘probefiltering’ functions available in the ‘DNAMarray’ R-package were used for further processing of array data. Finally, 412,915 probes contained on the EPIC array, were used for further analysis. Most significantly differentially methylated CpGs between disease entities were identified by fitting a regression model with the disease as the target variable using the ‘limma’ R package. All pairwise comparisons between disease groups were identified as contrasts and included in the analysis. No surrogate variable adjustments (‘sva’ R package) or batch corrections were necessary. After identification of 108 unique most significantly differentially methylated CpGs, unsupervised dimensionality reduction for cluster analysis was performed. Uniform Manifold Approximation and Projection (UMAP) for general non-linear dimensionality reduction was used for visualization [45]. After identification of disease clusters, care was taken that no cluster was confounded or correlated with any other variable such as sex, age at onset, age at surgery, duration of epilepsy, lobe (**Supplement Fig. 1**). Subsequently, additional hierarchical cluster analysis was performed.

### Machine and deep learning

’Scikit-learn,’ ‘imbalanced-learn,’ ‘Keras,’ and ‘Tensorflow’ were used as Python-based packages to leverage machine and deep learning. The processed methylation data obtained was split into a training and independent test set. Care was taken that disease classes were stratified across the sets evenly. Afterwards, the training and test set were upsampled to retrieve sample groups of almost the same sizes using the Synthetic Minority Oversampling Technique (SMOTE) of the ‘imbalanced-learn’ package. Various ‘classic’ machine learning algorithms (using ‘Scikit-learn‘) were spot-checked on their performance on the data via 10-fold cross-validation (**Supplement Fig. 1**). Independently, six-fold stratified cross-validation was performed using a deep neuronal net architecture. The network consisted of four fully connected layers (one input and one output layer). The first three fully connected layers used a rectified linear activation function followed by a batch normalization layer and a subsequent dropout layer. Only the last layer had a sigmoid activation function. Neither batch normalization nor dropout did follow. The training was performed using a batch size of 64, cycling the learning rate between 0.00001 and 0.09 on every epoch for a total of 66 epochs, which was identified by early stopping. All parameters described were evaluated via a prior grid search. The network and training process was implemented using the Keras and Tensorflow Python-based deep learning frameworks.

### FISH and RNA sequencing

FISH analysis was performed on FFPE-sections from ten representative cases with and without a change in copy number profile. Sections from lesional (with PMG) and perilesional tissue blocks (no PMG) of the same patients were analyzed. Two color interphase FISH analysis was performed using 1q21 CKS1B Spectrum Orange / 1p32 CDKN2C Spectrum Green FISH Probe Kit (Vysis; Abbott). Pre-treatment of slides, hybridization, post-hybridization processing, and signal detection were performed as reported elsewhere [15].Samples showing sufficient FISH efficiency (>90% nuclei with signals) were evaluated by two independent investigators. Signals were scored in at least 300 non-overlapping, intact nuclei. Trisomy/gain of 1q was defined as >15 % of nuclei containing three or more signals for the respective locus probe, if no such findings were detected for the 1p locus (to rule out polyploidy).

RNA was obtained from the same samples and, upon reverse transcription, was subjected to next-generation sequencing on a NextSeq 500 (Illumina, San Diego, CA, USA), as described previously [61]. We used deFuse [46] and Arriba (https://github.com/suhrig/arriba/) methods for the detection of gene fusions.

### Data Availability

The methylation data was deposited in NCBIs Gene Expression Omnibus (GEO, http://www.ncbi.nlm.nih.gov/geo) and will be made accessible upon publication (accession number XXX). **Supplement Table 1** includes IDAT-file names for assignment to patient characteristics.

## Results

### Novel methylation cluster defines PMG among other MCD

To determine, whether DNA methylation signatures can be used to classify structurally related MCD molecularly, we used the methylation data from surgical brain samples obtained from 26 pharmacoresistant epilepsy patients with a histopathological diagnosis of PMG (**Table 1**) and 62 MCD and no-MCD reference cases (i.e., FCD type 2a, FCD type 2b, HME, TLE) as well as from no-seizure autopsy controls (CTRL, microdissected white matter and neocortex; **Supplement Table 1**). We performed unsupervised dimensionality reduction and hierarchical cluster analysis. In addition to previously described FCD and TLE specific methylation classes [36], PMG in our analysis formed a novel separate cluster in the UMAP dimensionality reduction (**Fig. 1a**). No confounding correlation with any other variable of our data was detected (e.g., sex, age, lobe; **Supplement Fig. 1**). Hierarchical cluster analysis confirmed the separation of samples at the disease level (**Fig. 1b**).

**Figure 1:**
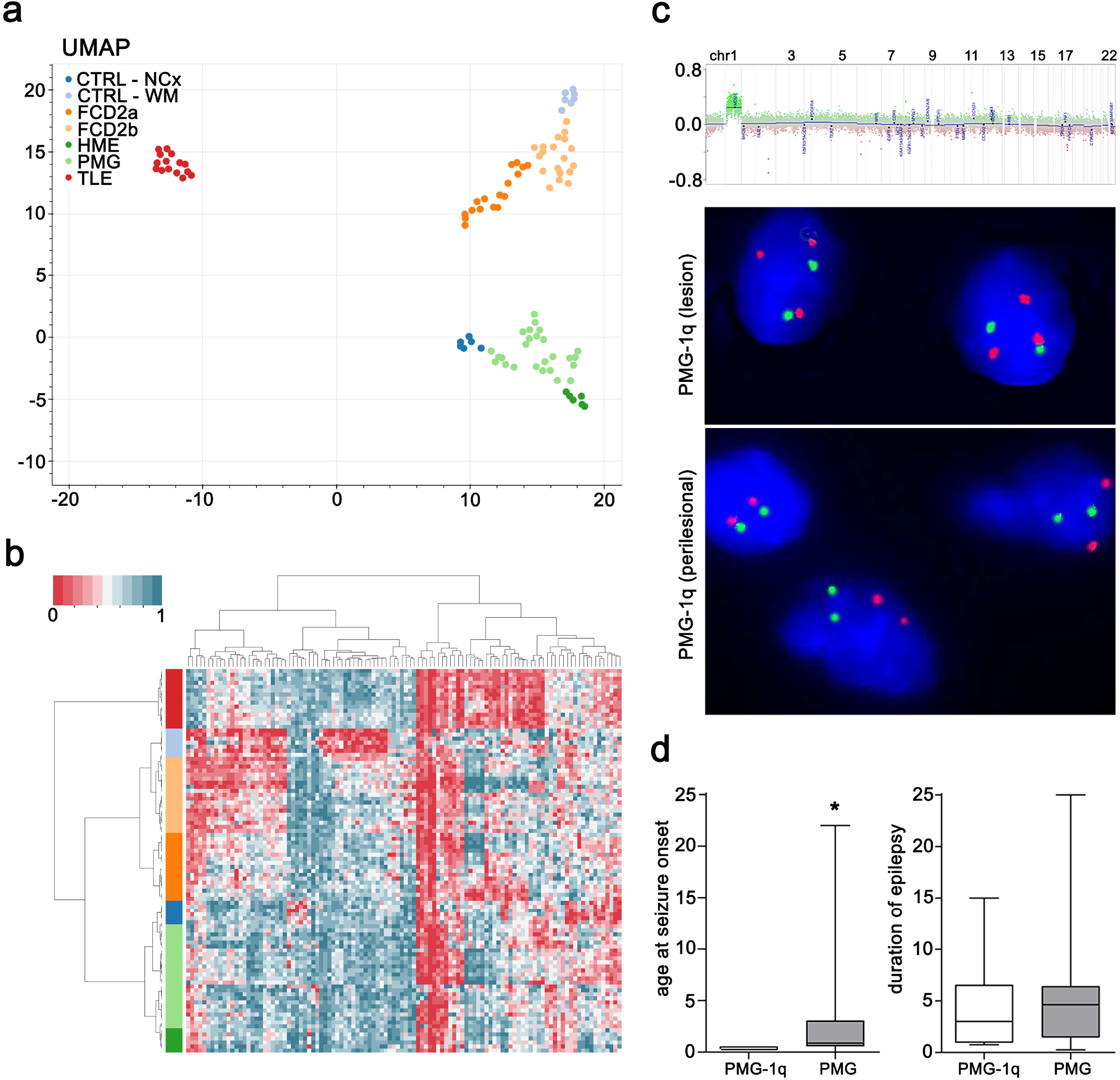
**(a)** UMAP and **(b)** hierarchical cluster analysis of PMG, reference MCD, non-MCD epilepsy, and no-seizure autopsy controls. Twenty-seven cases with histological diagnosis of PMG are indicated in light green. PMG cases formed a novel distinct methylation group. **(c)** Copy number profiling analysis indicating duplication of chromosome arm 1q and FISH confirming brain somatic 1q triploid nuclei in the center of the lesion, but not in adjacent, architecturally normal-appearing tissue of the same patient. **(d)** 1q duplication in PMG patients associated with significantly earlier onset of seizures, but not longer duration of epilepsy before surgery. Chr – chromosome; CTRL – control; NCx – neocortex; WM – white matter; FCD – focal cortical dysplasia; HME – hemimegalencephaly; PMG – polymicrogyria; TLE – temporal lobe epilepsy; UMAP - uniform manifold approximation and projection

### Brain mosaic 1q triploidy as a new defining feature of some PMG

Next we performed copy number profile analysis from DNA methylation data, which identified a uniform duplication spanning the entire long arm of chromosome 1 in 7/26 PMG cases (**Fig. 1c**). FISH analysis identified a mosaic distribution of about 50% 1q triploid nuclei in the center of the PMG lesion of respective cases (mean±SEM = 51.2 % ± 1.3), but not in adjacent, architecturally normal-appearing tissue (mean±SEM = 7.2 % ± 3.0; **Fig. 1c**, **lower panel**). Due to the retrospective study design, no blood samples were available from patients to confirm the mosaicism of the 1q duplication further. RNA-Seq analysis from same samples revealed the absence of translocations and gene fusions in all PMG patients with and without 1q copy number variation.

Our findings provide evidence for a previously unrecognized PMG specific methylation class and a brain mosaic chromosomal rearrangement affecting 1q in a subgroup of PMG patients.

### 1q trisomy associated with focal or hemispheric PMG, absence of hemimegalencephaly, early-onset severe focal epilepsy and developmental delay

On MRI, PMG cases with 1q gain presented as unilateral focally restricted frontal or hemispheric PMG (**Fig. 2a**, **Supplement Fig. 2**). Histomorphological features of the PMG lesion included abnormally folded sulci without pial opening. The cortical ribbon was small and mostly only 4-layered (**Fig. 2b**). One case showed protoplasmic astrocytic inclusions (**Fig. 2c**, blue arrow). All 1q positive PMG cases were clinically associated with a significantly earlier onset of seizures in the first few months of life (mean onset_PMG1q_ = 0.4 ± 0.04 years, n=7) as compared to the other PMG in our cohort (mean onset_PMG_ = 2.3 ± 0.7 years, n=19, Mann-Whitney test, p=0.018; **Fig. 1d)**. The mean duration of epilepsy before surgery was not different between the two molecular PMG subgroups (mean duration_PMG1q_ = 4.3 ± 1.9 years, n=7; mean duration_PMG_ = 5.5 ± 1.4 years, n=19; Mann-Whitney test, p=0.48; **Fig. 1d**). Based on the available routine clinical reports there was no evidence for a specific seizure pattern that would distinguish this patient group. Six out of seven PMG patients with brain mosaic 1q duplication presented with a severe combinatorial developmental delay including cognition, speech, and motor. Five out of seven patients had a post-surgical outcome of Engel class 1 five years after surgery [19]. Two patients did not become seizure-free due to incomplete resection of the epileptogenic zone to avoid hemiparesis.

**Figure 2:**
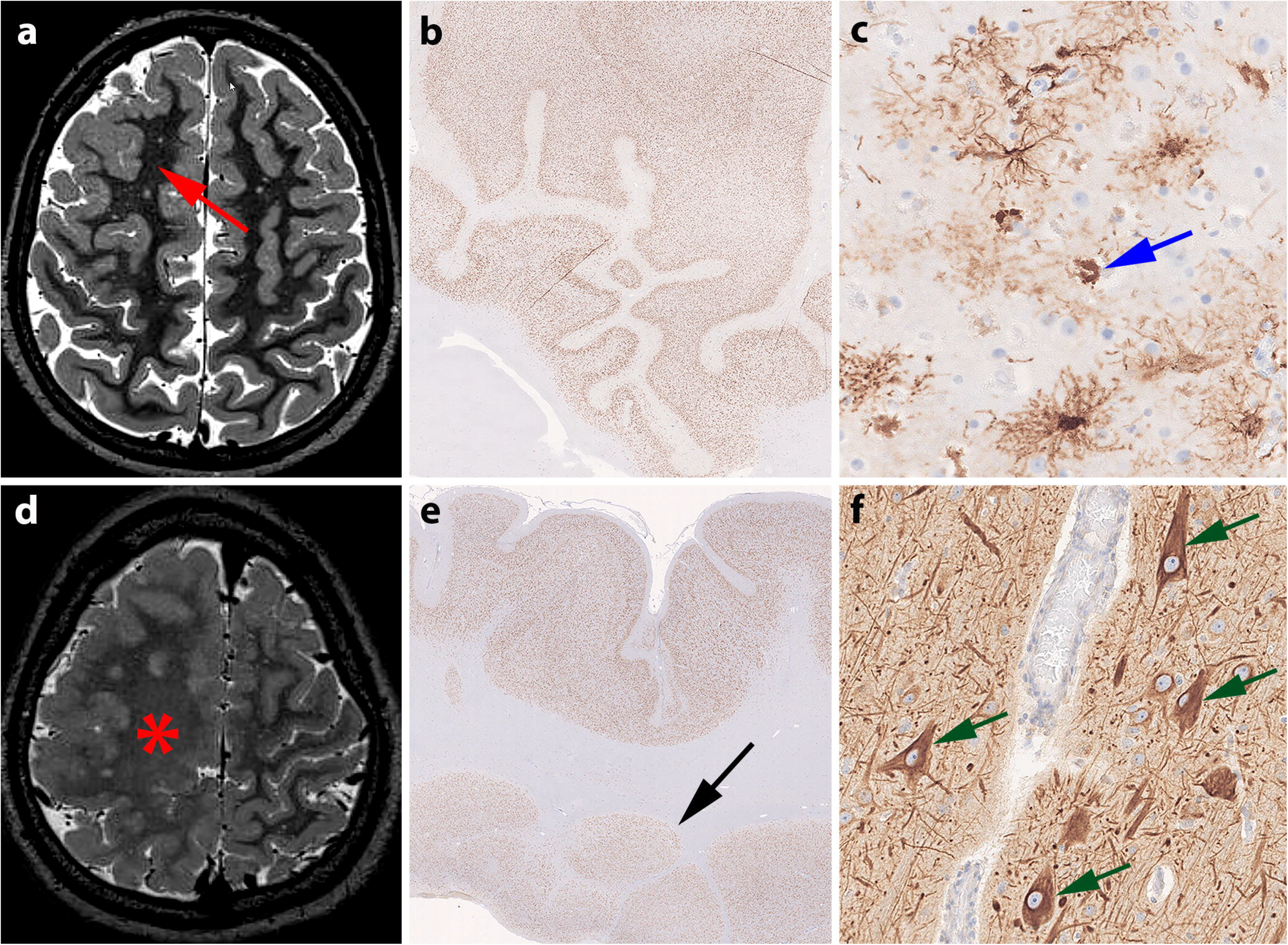
MRI and histology of representative PMG cases with and without brain mosaic 1q gain. **(a)** 15-year old male patient (#6) with seizure onset at 3 months and an MRI-positive lesion in the right frontal lobe with cortical thickening and no hyperintense T2/FLAIR signal and no transmantle sign (arrow). **(b)** NeuN staining revealed abnormally folded sulci without pial opening. The cortical ribbon is small and mostly 4-layered. **(c)** GFAP-immunohistochemistry revealed protoplasmic astrocytic inclusions (blue arrow [27]). **(d)** 17-year old male patient (#18) with seizure onset at 8 months and right hemimegalencephaly (asterisk). (**e**) neocortical ribbon with small sulci without pial opening and nodular heterotopias in the white matter (black arrow; NeuN immunochemistry). **(f)** Dysmorphic neurons accumulating non-phosphorylated neurofilament protein (green arrows, SMI32-immunohistochemistry)

In contrast, PMG cases without 1q duplication showed greater variability in their clinical presentation. In addition to the significantly later epilepsy onset, these cases more frequently presented with their PMG as part of a hemimegalencephaly (**Fig. 2d**, red asterisk) or otherwise complex cortical malformation with, e.g., cortical thickening, broad signal alterations in the white matter, blurring of the grey-white matter junction, periventricular or subcortical nodular heterotopias (**Fig. 2e**, black arrow), and FCD type 2 (**Fig. 2f**, green arrows), schizencephaly or microcephaly (**Table 1**). Two patients in this subgroup were diagnosed with Sturge-Weber-Syndrom. Five patients had no or only mild-to-moderate intellectual disability and developmental delay. Thirteen patients were reported with severe combinatorial developmental delay. For one case we had no detailed information. One patient had a reported 22q11 microdeletion. Another one had a reported congenital cytomegalovirus infection. However, no systematic genetic or viral testing was performed in the cohort. Thirteen out of 19 patients became seizure free in this subgroup.

## Discussion

A growing list of PMG types and syndromes have been observed, most of which have not been completely delineated. Their description is primarily based on their localization (by MRI) and correlation with clinical aspects including developmental course, growth anomalies, and dysmorphism, seizure history, family history, and genetic testing of blood for PMG associated genes [64]. We studied a series of 26 patients with pharmacoresistant epilepsy that underwent surgical treatment and were diagnosed with a PMG based on imaging and histology. From this cohort, we extracted a set of 7 patients exhibiting a unique molecular fingerprint along with specific clinical features including early-onset epilepsy in the first months of life and severe combinatorial developmental delay, thereby defining a distinct PMG entity. The molecular fingerprint relied on both a highly characteristic methylation profile and invariable detection of a brain mosaic duplication of the long arm of chromosome 1.

MCD are the most frequent causes of childhood epilepsies, carrying a lifelong disability perspective and reduced quality of life [12]. They represent a wide range of cortical lesions resulting from derangements of normal intrauterine developmental processes and involving cells implicated in the formation of the cortical mantle [59]. The pathological features depend mainly on the timing of the defect in the developmental processes and the cause, e.g., abnormal proliferation or apoptosis, differentiation, neuronal migration, or layering [63]. Classification of MCD has proven challenging as two or more forms of MCD may coexist. Also, a particular defect in corticogenesis may give rise to more than one morphological subcategory of MCD, and, conversely, a morphological subtype of MCD may have more than one mechanism for its formation [2]. Thus, there is a need to complement current classification of MCD with molecular-genetic data.

PMG are a heterogeneous group of MCD characterized by an excessive number of abnormally small gyri and abnormal cortical lamination. They can vary in size and localization, be unilateral or bilateral symmetrical or asymmetrical, and further occur as an isolated malformation or accompany other brain malformations such as HME, microcephaly, shizencephaly, periventricular nodular heterotopia, and FCD among others. As the classification of PMG remains challenging to date if solely based on imaging and histopathology, we tested whether PMG were associated with a distinct molecular fingerprint.

The impact of integrating molecular genotypes with histopathological phenotypes for the classification of brain tumors was stated by the WHO in 2016 [44]. Beyond disease-related genetic variants, genomic DNA methylation classifiers have been identified as a valuable source in the decision-making process for disease diagnosis, prognosis, and treatment [16]. Recently we showed that genomic DNA methylation signatures also distinguished major FCD subtypes from TLE patients and non-epileptic controls [36]. Here, we confirmed and extended this finding in a new independent patient cohort, including FCD among other MCD (i.e., PMG, HME), TLE, and no seizure autopsy controls. All disease groups were identified to fall into distinct methylation classes, as shown by unsupervised dimensionality reduction and hierarchical cluster analysis. At the point of writing this manuscript, there had been no previous report on PMG-specific DNA methylation signatures.

CNVs, i.e., deletions or duplications of a stretch of chromosomal DNA, can be part of normal genomic variation, but also cause or increase the risk for disease [47]. In the present study, we identified a somatic duplication of the entire long arm of chromosome 1 in surgical brain tissue obtained from PMG patients. Chromosome 1 is the largest human chromosome, with a length of 248,956,422 bp and 2,055 coding and 2,092 non-coding genes (ENSEMBL, GRCh38.p13). It is highly susceptible to genetic variations such as polymorphisms or mutations, and many diseases have been linked to these abnormalities. Complete monosomy is invariably lethal before birth and complete trisomy is lethal within days after conception [5]. Symptoms of congenital partial deletions and partial duplications generally depend on the size and location of the anomaly, and the genes involved. Several 1q related microdeletion and duplication syndromes or translocations are known. Features that may occur in respective patients include developmental delay and learning disabilities, slow growth and short stature, various congenital disabilities (such as cleft palate or a heart defect), specific facial features (such as a small, receding jaw), as well as neoplasias. This manifold of diseases highlights the central role of genes on the long arm of chromosome 1 in overall development, but also brain structure and function. For example, recurrent rearrangements of 1q21.1 are associated with microcephaly or macrocephaly, developmental, behavioral, and psychiatric problems (e.g., autism spectrum disorders, attention-deficit disorder, learning disabilities, schizophrenia), and seizures [14, 24, 48, 52]. 1q24 deletions cause a phenotype of intellectual disability, growth retardation, microcephaly, and facial dysmorphism [17]. Patients with 1q41 microdeletion syndrome were reported to present with seizures, mental retardation or developmental delay, and dysmorphic features at varying degrees [58]. Another microdeletion affecting 1q43q44 was associated with corpus callosum abnormalities, microcephaly, intellectual disability, and seizures [13, 22, 28, 42]. Of the large variety of germline mutations that have been previously associated with PMG, two map to chromosome 1q: *AKT3* (chr1q43-44) and *FH* (chr1q43), [20, 26, 55]. Of all CNVs that have been described in PMG [23, 56], none mapped to 1q before. To the best of our knowledge this is the first description of a 1q duplication syndrome affecting the entire long arm of chromosome 1 in focal epilepsy patients with a histopathological diagnosis of PMG.

Somatic variants are the result of postzygotic DNA mutational events that lead to clonal cell populations in an organism [43]. They are not inherited from a parent and not transmitted to the offspring (with the exception of germline mosaicism). The variant allele frequency detected in the brain specimens depends on multiple factors: the timing of occurrence during neurodevelopment, the impact on cellular proliferation, and the presence of selective pressure on the mutated cells. In the developing fetal brain, the cellular proliferation rate of cortical precursors between the 4th and 24th week of gestation is higher than in any other organ at any developmental stage [29]: Thus, somatic variants including single nucleotide variants and small insertions/deletions, CNVs or whole chromosome gains or losses, and mobile element insertions are a likely event in neuronal and glial cell lineages. Somatic variants occurring in mitotic cells lead to the development of a clone of mutated cells, that can represent a major part of an organism, such as a complete tissue layer, or just a small percentage of cells within an organ, and may or may not be visible [9]. Here we described a lesion-associated duplication of 1q in a subset of PMG patients with severe developmental and epileptic encephalopathy and very early onset of seizures in the first few months of life. We identified a mosaic distribution of 1q triploid nuclei in the center of the PMG lesion, but not in peri-lesional normal-appearing tissue. Due to the retrospective study design, no blood samples were available from our PMG patients to further verify mosaicism. There have been no previous descriptions of brain somatic variants in PMG. So far, somatic variants in focal epilepsy patients with an underlying MCD have been identified only in FCD type 2 and HME [1, 3, 4, 20, 41, 49–51].

Illumina Infinium DNA Methylation EPIC Bead Chip Arrays are an established tool for molecular classification of brain tumors and other cancers [16]. Here we tested, whether integrating clinico-pathological and epigenetic analysis is also helpful to characterize and classify epileptogenic cortical malformations. We previously used Methyl-capture massive parallel sequencing in fresh frozen surgical tissue and showed that DNA methylation signatures i) can distinguish epileptic from non-epileptic healthy control tissue [21, 35, 36]; and ii) are indicative of the underlying etiology and histopathology [21, 36]. We now confirmed and extended these findings in a new independent sample cohort using FFPE tissue and a different, cost-efficient, and clinically widely distributed profiling platform technology. Our data show that somatic 1q trisomy associated PMG cases and PMG without this chromosomal imbalance formed distinct methylation classes. Differential methylation affected not only regions on the long arm of chromosome 1, but was distributed on other autosomes (data not shown) suggesting that DNA methylation changes were not only linked to the 1q duplication, but may be an integral part of the epileptogenic process, including the development of a structural lesion, seizures, and co-morbidities [11, 33, 34, 38]. Future studies will need to address the potential causative role of DNA methylation changes in the development of cortical malformations and associated seizures. We have not performed any functional pathway analysis as we have shown earlier the superiority of high throughput sequencing over arrays for identification of genomic DNA methylation changes in an unbiased manner and sufficient coverage, thereby providing new insights into molecular disease pathogenesis [36].

Seventeen PMG patients in the present study, 5 with 1q duplication and 13 without, had a post-surgical outcome of Engel class 1 five years after surgery. Irrespective of the small study cohort, our data suggest that surgical treatment is recommended in pharmacoresistant focal epilepsy patients with unilateral PMG and associated with favourable seizure outcome.

## Conclusion

We describe the invariable coincidence of a specific methylation profile with the presence of a brain somatic trisomy involving chromosome 1q in a distinct group of PMG patients without hemimegalencephaly, with early-onset epilepsy in the first months of life and severe developmental delay. Other PMG showed no chromosomal imbalances, formed a distinct methylation class in hierarchical cluster analysis, more often presented as complex malformation on MRI and histologically, and showed considerable higher variability of clinical phenotype. Our data support an integrated molecular-genetic and histological disease classification of cortical malformations including PMG. Moreover, we identify a new patient group with unilateral PMG, very early-onset focal epilepsy and brain mosaic 1q whole arm duplication.

## Acknowledgments

We kindly thank B. Rings for her expert technical assistance. Our work was supported by the European Union’s Seventh Framework Program (DESIRE project, grant agreement 602531) and European Reference Network EpiCare (grant agreement 769501). A. Korshunov is supported by the German Helmholtz Association Research Grant (project number HRSF-0005). A. Mühlebner is supported by the Dutch Epilepsy Foundation (project number 20-02).

## Supplements

**Supplement Figure 1**

**Legend to Supplement Fig. 1: (a)** A neural network was trained for DNA methylation-based disease classification. **(b-c)** Comparison with other machine learning methods showed superior performance of the neural network in 6-fold cross-validation. **(d)** UMAP plots were generated from 108 most significantly differentially methylated positions with potentially confounding variable influence. Samples were labelled with the covariates age at onset, age at surgery, duration of epilepsy, sampled lobe (temporal, frontal, parietal, insular, NOS – not otherwise specified), cortex (green) or white matter (red), and sex, male (green), female (red). Clustering of samples was not driven by the tested covariates.

**Supplement Figure 2**

**Legend to Supplement Figure 2:** Representative MRIs of PMG-1q patients. No HME was present in this patient subgroup.

**Supplement Figure 3**

**Legend to Supplement Figure 3:** Representative MRI of PMG patients without 1q duplication. In these cases, PMG was more frequently part of HME or other complex malformations.

**Supplement Table 1**

**Legend to Supplement Table 1:** Clinical summary of reference cohort and idat allocation.

