## Supplementary figures and images for "Mosaic trisomy of chromosome 1q in human brain tissue associates with unilateral polymicrogyria, very early-onset focal epilepsy, and severe developmental delay"

### Supplemental Figure 1

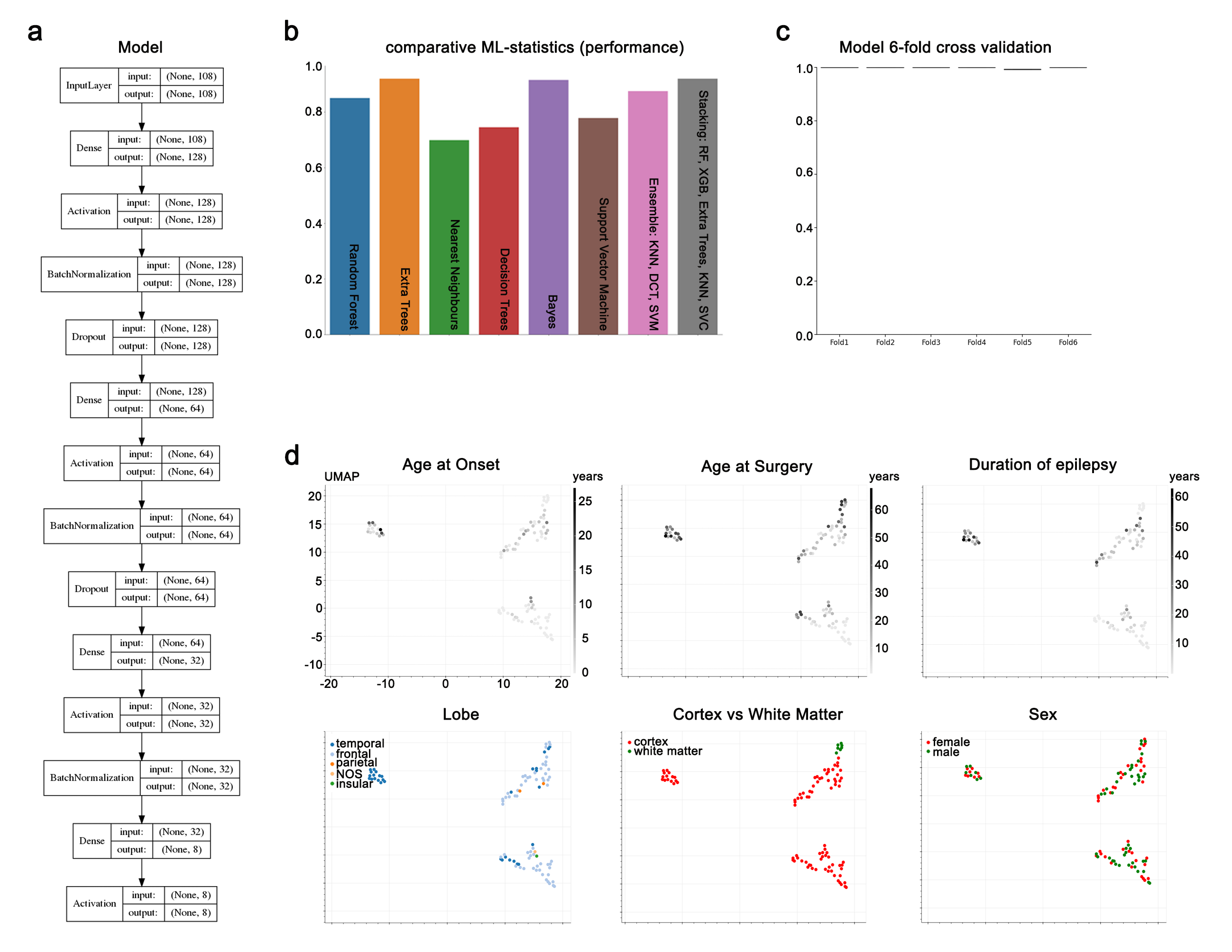

### Supplemental Figure 2

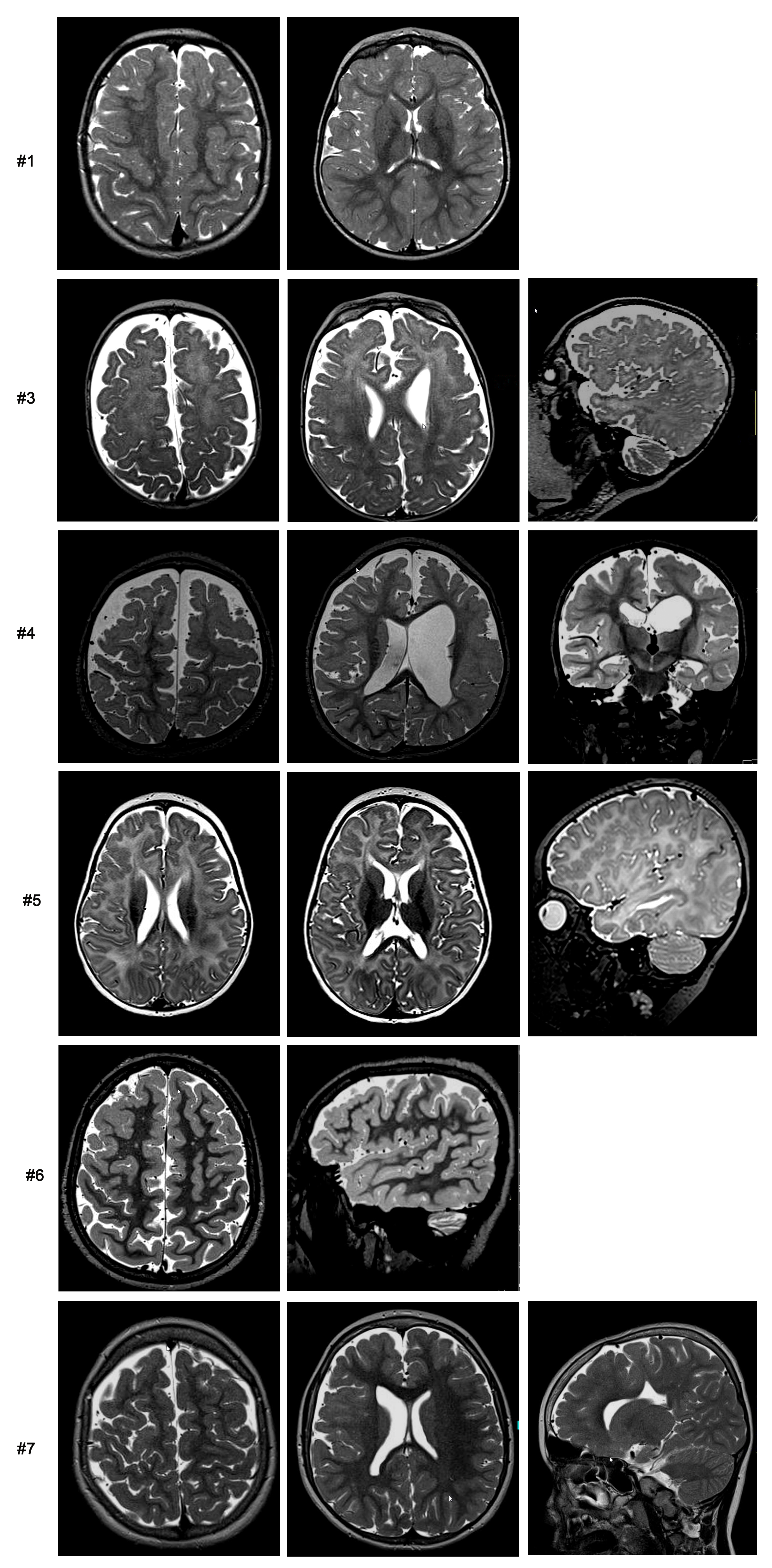

### Supplemental Figure 3

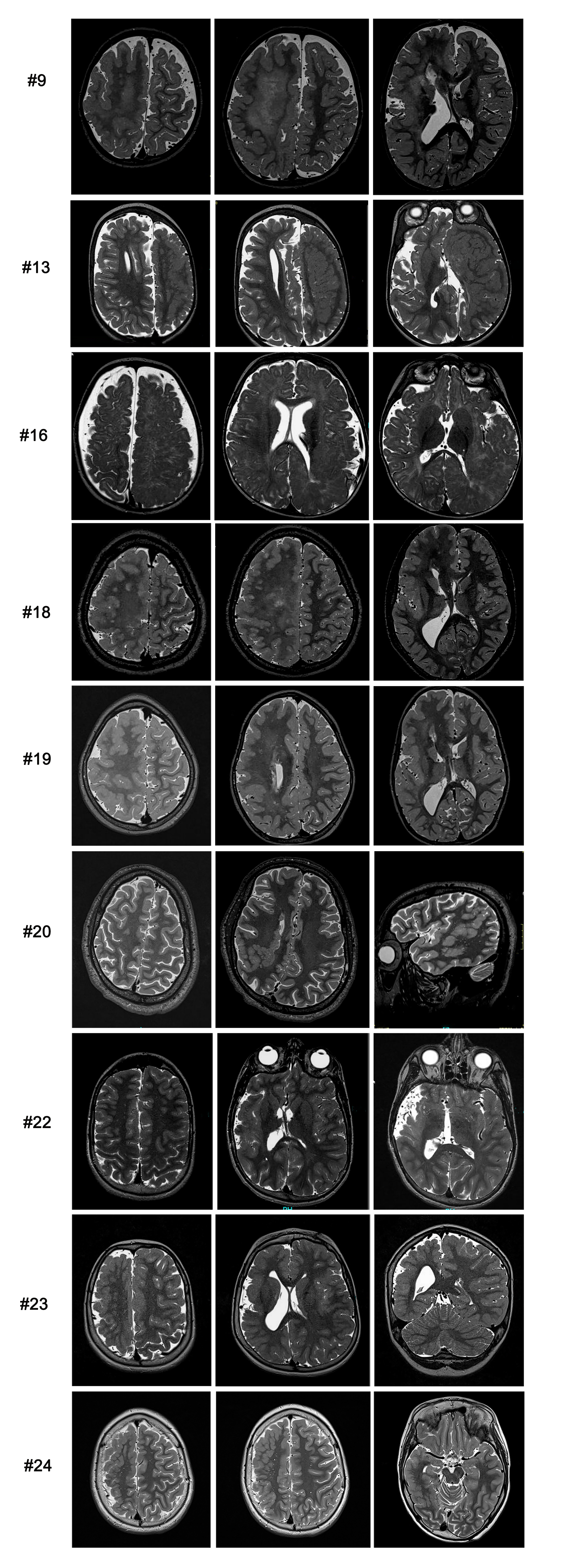
