## Supplemental Table 1 for "Mosaic trisomy of chromosome 1q in human brain tissue associates with unilateral polymicrogyria, very early-onset focal epilepsy, and severe developmental delay"

| # | pathology | lobe | sex | age_onset | duration | age_surgery |
|---|---|---|---|---|---|---|
| 1 | PMG | frontal | F | 0 | 3 | 3 |
| 2 | PMG | frontal | M | 0 | 1 | 1 |
| 3 | PMG | frontal | M | 0 | 1 | 1 |
| 4 | PMG | frontal | M | 0 | 3 | 3 |
| 5 | PMG | frontal | F | 0 | 1 | 1 |
| 6 | PMG | NOS | F | 0 | 15 | 15 |
| 7 | PMG | frontal | F | 0 | 7 | 7 |
| 8 | PMG | temporal | M | 2 | 6 | 8 |
| 9 | PMG | frontal | M | 0 | 3 | 3 |
| 10 | PMG | frontal | M | 1 | 1 | 2 |
| 11 | PMG | temporal | F | 11 | 6 | 17 |
| 12 | PMG | frontal | M | 0 | 6 | 6 |
| 13 | PMG | frontal | M | 1 | 2 | 3 |
| 14 | PMG | frontal | F | 0 | 1 | 2 |
| 15 | PMG | frontal | F | 0 | 2 | 2 |
| 16 | PMG | frontal | M | 0 | 1 | 1 |
| 17 | PMG | frontal | M | 1 | 7 | 8 |
| 18 | PMG | frontal | M | 1 | 16 | 17 |
| 19 | PMG | frontal | F | 0 | 5 | 5 |
| 20 | PMG | frontal | M | 10 | 25 | 35 |
| 21 | PMG | temporal | M | 3 | 2 | 5 |
| 22 | PMG | temporal | M | 3 | 10 | 13 |
| 23 | PMG | frontal | M | 2 | 5 | 7 |
| 24 | PMG | Insel | M | 3 | 6 | 9 |
| 25 | PMG | frontal | F | 4 | 4 | 8 |
| 26 | PMG | frontal | M | 0 | 2 | 2 |
| 27 | CTRL - NCx | temporal | F | | | 11 |
| 28 | CTRL - NCx | temporal | M | | | 27 |
| 29 | CTRL - NCx | frontal | M | | | 27 |
| 30 | CTRL - NCx | temporal | F | | | 13 |
| 31 | CTRL - NCx | temporal | F | | | 49 |
| 32 | CTRL - NCx | frontal | F | | | 49 |
| 33 | CTRL - WM | temporal | F | | | 11 |
| 34 | CTRL - WM | frontal | M | | | 27 |
| 35 | CTRL - WM | temporal | F | | | 13 |
| 36 | CTRL - WM | temporal | F | | | 49 |
| 37 | CTRL - WM | frontal | F | | | 49 |
| 38 | CTRL - WM | frontal | M | | | 52 |
| 39 | CTRL - WM | frontal | M | | | 52 |
| 40 | FCD 2A | frontal | M | 2 | 2 | 4 |
| 41 | FCD 2A | frontal | M | 0 | 1 | 1 |
| 42 | FCD 2A | frontal | F | 0 | 3 | 3 |
| 43 | FCD 2A | frontal | F | 8 | 0 | 8 |
| 44 | FCD 2A | frontal | F | 1 | 9 | 9 |
| 45 | FCD 2A | frontal | M | 4 | 9 | 13 |
| 46 | FCD 2A | frontal | F | 3 | 7 | 10 |
| 47 | FCD 2A | temporal | M | 0 | 6 | 6 |
| 48 | FCD 2A | frontal | F | 10 | 8 | 18 |
| 49 | FCD 2A | frontal | M | 5 | 6 | 11 |
| 50 | FCD 2A | frontal | M | 3 | 14 | 17 |
| 51 | FCD 2A | frontal | M | 9 | 15 | 24 |
| 52 | FCD 2A | frontal | F | 0 | 19 | 19 |
| 53 | FCD 2A | frontal | F | 4 | 11 | 15 |
| 54 | FCD 2A | parietal | F | 1 | 11 | 12 |

| | | | | | | |
|---|---|---|---|---|---|---|
| 55 | FCD 2A | frontal | F | 0 | 45 | 45 |
| 56 | FCD 2A | frontal | F | 2 | 30 | 32 |
| 57 | FCD 2B | frontal | M | 1 | 1 | 2 |
| 58 | FCD 2B | frontal | F | 0 | 3 | 3 |
| 59 | FCD 2B | parietal | M | 0 | 11 | 11 |
| 60 | FCD 2B | frontal | M | 6 | 2 | 8 |
| 61 | FCD 2B | frontal | M | 2 | 5 | 7 |
| 62 | FCD 2B | temporal | M | 0 | 5 | 5 |
| 63 | FCD 2B | frontal | M | 4 | 2 | 6 |
| 64 | FCD 2B | temporal | M | 0 | 5 | 5 |
| 65 | FCD 2B | frontal | F | 3 | 7 | 10 |
| 66 | FCD 2B | frontal | M | 1 | 7 | 8 |
| 67 | FCD 2B | frontal | F | 1 | 15 | 16 |
| 68 | FCD 2B | frontal | F | 4 | 11 | 15 |
| 69 | FCD 2B | temporal | M | 1 | 15 | 16 |
| 70 | FCD 2B | frontal | F | 5 | 15 | 20 |
| 71 | FCD 2B | frontal | M | 5 | 15 | 20 |
| 72 | FCD 2B | frontal | F | 13 | 30 | 43 |
| 73 | FCD 2B | frontal | M | 3 | 40 | 43 |
| 74 | FCD 2B | frontal | M | 2 | 36 | 38 |
| 75 | FCD 2B | temporal | F | 5 | 41 | 46 |
| 76 | HME | frontal | M | 0 | 1 | 1 |
| 77 | HME | frontal | M | 1 | 1 | 1 |
| 78 | HME | frontal | F | 0 | 2 | 2 |
| 79 | HME | frontal | F | 0 | 1 | 1 |
| 80 | HME | frontal | F | 0 | 2 | 2 |
| 81 | HME | frontal | F | 0 | 1 | 1 |
| 82 | TLE | temporal | M | 16 | 10 | 26 |
| 83 | TLE | temporal | F | 2 | 17 | 19 |
| 84 | TLE | temporal | F | 27 | 16 | 43 |
| 85 | TLE | temporal | F | 4 | 11 | 15 |
| 86 | TLE | temporal | F | 1 | 15 | 15 |
| 87 | TLE | temporal | M | 6 | 23 | 29 |
| 88 | TLE | temporal | F | 18 | 23 | 41 |
| 89 | TLE | temporal | M | 6 | 44 | 50 |
| 90 | TLE | temporal | F | 3 | 25 | 28 |
| 91 | TLE | temporal | M | 1 | 44 | 45 |
| 92 | TLE | temporal | M | 15 | 25 | 40 |
| 93 | TLE | temporal | F | 5 | 28 | 33 |
| 94 | TLE | temporal | M | 1 | 47 | 48 |
| 95 | TLE | temporal | M | 1 | 54 | 55 |
| 96 | TLE | temporal | F | 5 | 63 | 68 |

| idat |
|---|
| 203220070058_R06C01 |
| 203220070058_R07C01 |
| 203219750116_R02C01 |
| 203219750116_R04C01 |
| 203219730055_R03C01 |
| 203219730055_R06C01 |
| 203511880025_R05C01 |
| 203220070058_R05C01 |
| 203219750116_R08C01 |
| 203219730055_R04C01 |
| 203219730055_R05C01 |
| 203511880014_R08C01 |
| 203511880025_R01C01 |
| 203511880025_R02C01 |
| 203511880025_R03C01 |
| 203511880025_R04C01 |
| 203511880025_R06C01 |
| 203219750146_R05C01 |
| 203511880026_R02C01 |
| 203511880026_R03C01 |
| 203220070058_R04C01 |
| 203220070058_R08C01 |
| 203219750116_R03C01 |
| 203219750116_R05C01 |
| 203219750116_R06C01 |
| 203219750116_R07C01 |
| 202818860053_R04C01 |
| 202818860053_R06C01 |
| 202818860053_R08C01 |
| 202931510124_R04C01 |
| 202931510124_R06C01 |
| 202931510124_R08C01 |
| 202818860053_R05C01 |
| 202931510124_R01C01 |
| 202931510124_R05C01 |
| 202931510124_R07C01 |
| 202939390010_R06C01 |
| 202939390010_R08C01 |
| 202944920003_R06C01 |
| 202818860117_R02C01 |
| 202818860117_R06C01 |
| 202827620174_R03C01 |
| 202827620174_R04C01 |
| 202093110113_R03C01 |
| 202822930036_R07C01 |
| 202818860117_R05C01 |
| 202827620173_R08C01 |
| 202827620174_R01C01 |
| 202827620174_R06C01 |
| 202093110108_R07C01 |
| 202093110113_R01C01 |
| 202093110113_R02C01 |
| 202818860117_R01C01 |
| 202818860117_R04C01 |

202093110108_R08C01
202822930036_R06C01
202831040055_R05C01
202831040055_R08C01
202831040056_R01C01
202831040056_R06C01
202831040056_R08C01
202822930161_R03C01
202822930161_R05C01
202822930161_R06C01
202831040055_R01C01
202831040056_R04C01
202831040055_R03C01
202831040055_R07C01
202831040056_R05C01
202831040056_R07C01
202822930161_R02C01
202148010052_R02C01
202831040055_R04C01
202831040056_R03C01
202822930161_R04C01
202827620174_R07C01
202827620174_R08C01
203219750146_R04C01
203219750146_R06C01
203219750057_R01C01
203219750057_R02C01
202148010059_R02C01
202148010053_R08C01
202148010058_R02C01
202148010058_R03C01
202148010058_R06C01
202148010053_R06C01
202148010053_R07C01
202148010058_R01C01
202148010058_R04C01
202148010058_R07C01
202148010059_R01C01
202148010059_R03C01
202148010059_R04C01
202148010058_R05C01
202148010058_R08C01
